# Slower transitions between control states lead to reductions in cognitive flexibility over the lifespan

**DOI:** 10.1101/2025.08.27.672689

**Authors:** Ivan Grahek, Xiamin Leng, Alexander Fengler, Amitai Shenhav

## Abstract

Declines in cognitive flexibility are a hallmark of cognitive aging, but their causes remain elusive. Here, we examine a previously untested source of aging-related cognitive inflexibility, building on a dynamical systems framework for flexible cognition. We propose that cognitive inflexibility can arise in part from slower transitions through the space of distinct configurations of cognitive control needed to pursue different goals. To test this model, we had participants across the lifespan (aged 19-88) perform a cognitive task under different performance goals, each of which induced a different configuration of cognitive control. Using computational modeling, we measured dynamic changes in two control signals (attentional focus and response caution) as participants pursued distinct goals. This allowed us to test three potential sources of age-related decreases in cognitive flexibility: 1) diminished control capacity in environments that require more goal switching; 2) diminished range of control adjustments; and 3) slower transitions between control configurations. Of these, we found that age was only associated with transition speed. When given sufficient time to maintain a given goal, older adults were able to adjust control to a similar extent as younger adults. However, older adults were more likely to undershoot their target control configuration when frequently switching between goals, consistent with longer transition times between configurations. Our findings demonstrate the critical role that cognitive dynamics play in explaining the mechanisms through which cognitive inflexibility arises in older adulthood.

**Significance:** Meeting the shifting demands of daily life requires a person to constantly adjust the way in which they allocate their cognitive resources. As people get older, it becomes increasingly difficult to achieve the same levels of cognitive flexibility. We show that these deficits can be explained by age-related changes in the dynamics governing adjustments between cognitive states. Older adults take longer to adapt their cognitive state to a new goal, despite demonstrating preserved ability to reach desired states when given sufficient time. These findings validate a new model of cognitive flexibility and its variability across individuals, including forms of inflexibility seen in healthy aging and across a range of psychiatric and neurological disorders.

## Introduction

Cognitive flexibility is critical for successful goal-directed behavior, yet this core cognitive function declines with age. Relative to younger adults, older adults tend to exhibit worse performance when switching between tasks (Cepeda et al., 2001; Mayr, 2001; Wasylyshyn et al., 2011; Chen & Hsieh, 2023; Radović et al., 2025), when recovering from task interruptions (Morales et al., 2022), and when adjusting how they perform a task following changes in their environment (Kosciessa et al., 2024). While it is known that the capacity for cognitive flexibility is underpinned by a process of adjusting cognitive control settings in line with one’s current goals (e.g., what information a person attends to), it remains unclear how this process is altered in aging to result in greater cognitive inflexibility (Paxton et al., 2008).

Here, we propose and test a novel mechanism underlying cognitive inflexibility in older adulthood. We begin by recognizing that different goals require different configurations of multidimensional control configurations. For example, a person requires minimal levels of attentional focus and response caution to read the news while waiting for a medical appointment. However, once they enter the doctor’s office, they will need a different configuration (e.g., high focus and caution). Thus, the ability to adjust cognitive control configurations to changes in one’s goals or environment is critical, and research shows that young adults do so readily (Egner & Hirsch, 2005; Forstmann et al., 2008; Danielmeier & Ullsperger, 2011; Parro et al., 2018; Frömer et al., 2021; Leng et al., 2021; Grahek et al., 2023). Leveraging a dynamical systems perspective on cognitive flexibility (Ueltzhöffer et al., 2015; Steyvers et al., 2019; Musslick & Bizyaeva, 2024; Grahek et al., 2025), we recently proposed that adjusting control states requires gradual transitions in the space of possible control configurations, from the current state (e.g., low level of focus) to the target state required by the new goal (e.g., high level of focus). Under this view, the speed of transitions between control configurations becomes a crucial factor limiting cognitive flexibility (Braun et al., 2015; Kim & Bassett, 2019; Musslick & Cohen, 2021; Holroyd, 2023; Grahek et al., 2025). This sets up the prediction that a major source of cognitive inflexibility in older adulthood arises from decreases in the speed at which a person adjusts their control configuration to changing goals.

To test this prediction, we combine a novel behavioral paradigm with computational models that allow us to precisely measure changes in cognitive control states (Grahek et al., 2022, 2025). Participants (19-88 years old) performed a cognitive control task (Stroop) under two performance goals (focus on speed vs. accuracy) that reliably induced different control configurations (high vs. low levels of attentional focus and caution). By fitting process models of the task to each participant’s behavior, we were able to estimate how long it took people to transition between control configurations when their goals change, and to examine how this *adjustment speed* changes with age. Our task design allowed us to compare age-related changes in this novel measure of cognitive inflexibility to potential age-related changes in two other measures of cognitive inflexibility: (1) decreases in levels of overall control in goal-varying vs. stable environments (which we term *switching inflexibility*; Kray & Lindenberger, 2000; Cepeda et al., 2001; Mayr, 2001; Ferguson et al., 2021), and (2) decreases in levels of control adjustment for speed vs. accuracy goals (which we term *configural inflexibility*; Theisen et al., 2021; von Krause et al., 2022). We find that transition speed significantly decreases with age, whereas our other measures of cognitive flexibility do not. Thus, our findings demonstrate that aging is associated with changes in how people traverse the space of cognitive states, emphasizing the critical role of cognitive dynamics in understanding aging.

## Results

### Measuring the Speed of Control Adjustments

To study transitions between different control configurations, we had participants perform a control-demanding task (Stroop task) under different performance goals (Fig. 1A). Participants across a wide age range (N=245, 19-88 years old) were instructed to perform the task either as quickly as possible (speed goal) or as accurately as possible (accuracy goal). These goals each yield qualitatively distinct performance profiles, with participants performing the task faster and less accurately in the speed relative to the accuracy goal. Leveraging formal models of decision-making and cognitive control allows us to measure the distinct control configurations that underpin these performance profiles (Bogacz et al., 2010; Leng et al., 2021; Ritz et al., 2022).

**Figure 1.**
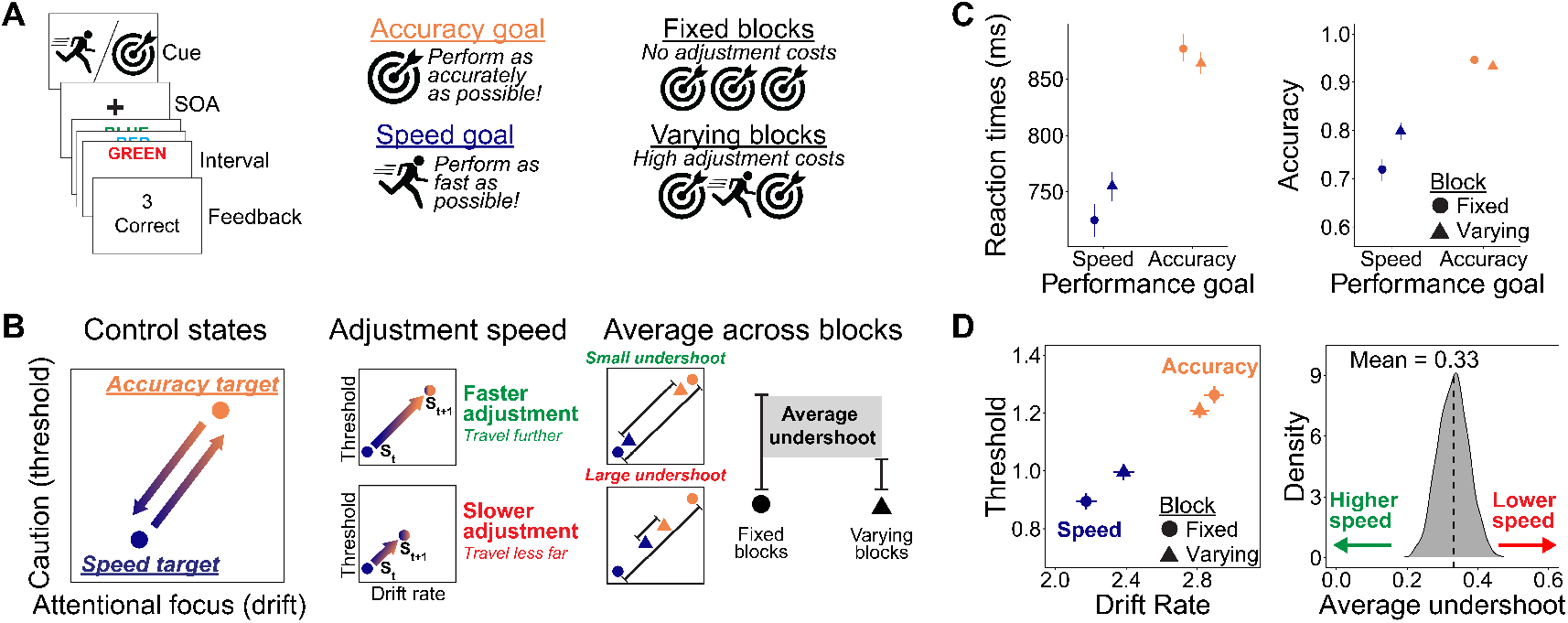
Behavioral task and measurement of transition speed. **A**. Participants performed a Stroop task (left) under two performance goals (speed vs. accuracy) signaled by the goal cue (middle). Participants completed the Stroop trial within limited time intervals (9-12s), completing as many trials as they wished in that time window. The task was structured into blocks (right), within the goal either always being the same (fixed) or with the goal randomly changing every interval (varying). **B**. The speed and accuracy performance goals require different target control configurations (left). Depending on the speed of control adjustments, participants will travel further or less far withing a unit of time (middle). Thus, blocks that require frequent control adjustments will lead to participants often undershooting their target state. In the aggregate, these undershoots are expected to result in smaller distances between control states in varying relative to fixed blocks (right). The transition speed determines the magnitude of this undershoot, with higher speeds leading to less undershoot because participants reach the target state faster. **C**. Participants are faster (left) and less accurate (right) when performing under the speed relative to the accuracy goal. The difference in performance between the two goals is smaller in varying relative to fixed blocks. Error bars represent 95% credible intervals. **D**. The two performance goals induce different control configurations inferred by fitting a Drift Diffusion model (left). Error bars represent 95% credible intervals. Euclidian distance between control configurations decreases in varying compared to fixed blocks (right). Posterior of the distance indicates that participants undershoot target configurations in varying blocks. There is substantial variability in the magnitude of the undershoot.

We have previously shown that whereas the accuracy goal yields high levels of attention and caution (which we will refer to as a participant’s *accuracy target*), the speed goal yields low levels of both (*speed target*; Fig. 1B-left) (Grahek et al., 2025). When participants are required to switch from one goal to another – e.g., from a speed target to an accuracy target or vice versa – they adjust the relevant control configurations gradually rather than all at once (Fig. 1B-middle). As a result, when these two goals are interleaved, participants tend to arrive at control configurations that fall short of their current target, for instance just below a high level of attention and caution when having switched from the speed goal to the accuracy goal. These shortfalls, in turn, provide an estimate of the time it takes for a person to transition between control states (Fig. 1B-right). Specifically, by comparing the distance between two control configurations in blocks with a single goal (fixed blocks) to blocks that require frequent switching between goals (varying blocks), we can ascertain how quickly or slowly they were able to traverse the distance between their target control configurations.

### People Gradually Transition Between Control Configurations

Across our whole sample, we find that participants are substantially slower (*b*=136.54; 95% CrI [117.74, 157.05]; *p*_*b*<0_=0.00) and more accurate (log odds: *b*=1.59; 95% CrI [1.41, 1.76]; *p*_*b*<0_=0.00; Fig. 1C and Table S1) under the Accuracy compared to the Speed goal. Fitting these behavioral data with a hierarchical drift diffusion model (Fengler et al., 2021; Wiecki et al., 2013; see Fig. S1 for posterior predictive checks), we replicate our finding that the differences between these two goals reflect differences in two model parameters: drift rate (reflecting the efficiency of information processing, an index of top-down attention) and decision threshold (an index of response caution). These parameter estimates revealed that the Speed and Accuracy goals engender substantially different control configurations (Fig. 1D-left and Table S1), with the Accuracy goal yielding higher drift rates (*b*=0.57; 95% CrI [0.50, 0.66]; *p*_*b*<0_=0.00) and higher thresholds (*b*=0.29; 95% CrI [0.25, 0.33]; *p*_*b*<0_=0.00) than the Speed goal.

Critically, as in our previous studies, we found that the performance profiles for the two goals were further apart from one another when performance goals were fixed for the entire block than when these goals were interleaved (RTs: *b*=44.11; 95% CrI [26.46, 62.73]; *p*_*b*<0_=0.00; accuracy: *b*=0.67; 95% CrI [0.50, 0.83]; *p*_*b*<0_=0.00; Fig. 1C and Table S1). In our model-based analyses, this manifest as an interaction whereby drift and threshold configurations were closer together on Varying relative to Fixed blocks (drift rate: *b*=0.29; 95% CrI [0.21, 0.38]; *p*_*b*<0_=0.00; threshold: *b*=0.16; 95% CrI [0.12, 0.20]; *p*_*b*<0_=0.28; Fig. 1D-left and Table S1).

Collectively, these findings are consistent with our model’s prediction (Figure 1B; Grahek et al., 2025) that people move gradually from one control configuration to another, therefore systematically undershooting each target configuration when forced to move between the two rather than remaining fixed on one goal for an extended period of time. Comparing the distance traversed between these two configurations on Fixed relative to Varying blocks thus provides a proxy for the time it takes to transition between these control configurations (Fig. 1B-right). Consistent with the interaction reported above, we find that this relative distance metric decreased substantially for Varying compared to Fixed blocks across our entire sample (*b*=0.33; 95% CrI [0.25, 0.42]; *p*_*b*<0_=0.00; Fig. 1D-right).

### Cognitive Inflexibility in Older Adults is Selectively Reflected in Control Adjustment Speed

With increasing age, people were on average significantly slower (*b*=223.11; 95% CrI [147.18, 306.71]; *p*_*b*<0_=0.00), while their accuracy did not change (*b*=0.11; 95% CrI [−0.53, 0.68]; *p*_*b*>0_=0.34; Table S1). These differences in task performance reflected age-related shifts in average control configuration towards significantly higher decision thresholds (*b*=0.19; 95% CrI [0.06, 0.33]; *p*_*b*<0_=0.00; cf. Theisen et al., 2021; von Krause et al., 2022), while drift rates did not reliably change (*b*=−0.11; 95% CrI [−0.52. 0.24]; *p*_*b*<0_=0.29; Table S1).

Our behavioral task and model-based analyses allow us to go beyond the overall age-related changes in control configurations and examine how adjustments of those configurations vary with age. In particular, we can test for age-related influences on three theoretical sources of cognitive inflexibility (Fig. 2): 1) the overall performance costs they incur when performing the task under frequent goal switches (goal switch flexibility); 2) the degree to which an individual is able to adjust control from one goal to another (configural flexibility); and 3) the speed with which they adjust control from one goal to another (adjustment speed, cf. Fig 1D). These forms of cognitive flexibility are not mutually exclusive, and thus our data can speak to whether one or more of them vary with age.

**Figure 2.**
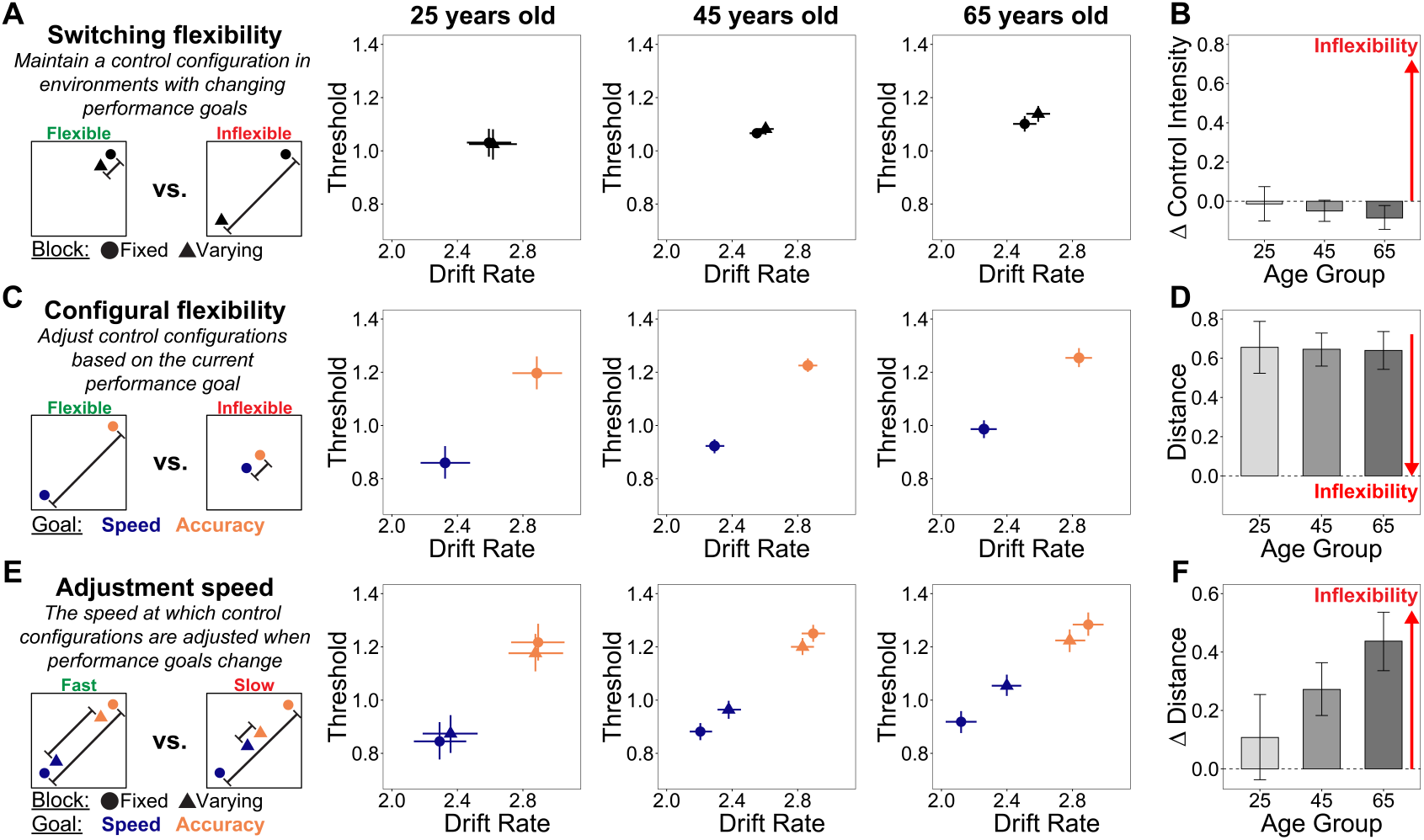
Changes in different types of cognitive flexibility across the life span. **A**. Left: Greater switching inflexibility would manifest as less control in blocks that require frequent goal switches (varying) relative to blocks that don’t require goal switches (fixed). Right: Average control configurations for fixed and varying blocks, separated by age bin. Error bars represent 95% credible intervals. **B**. A summary measure of the signed Euclidean distance between control configurations shows that switching flexibility does not decrease with age. Error bars represent 95% credible intervals. **C**. Greater configural inflexibility would manifest as smaller distances between control configurations under the two goals. Right: Average control configurations for speed and accuracy goals, separated by age bin. Error bars represent 95% credible intervals. **D**. A summary measure of the absolute Euclidean distance between speed vs. accuracy control configurations shows that configural flexibility does not decrease with age. Error bars represent 95% credible intervals. **E**. Greater adjustment speed would be reflected as greater reductions in the distance between Speed and Accuracy control configurations in varying vs. fixed blocks (due to greater undershooting; see Fig. 1D). **F**. With increasing age, transition speeds are lowered, and the amount of undershooting is increased. Error bars represent 95% credible intervals.

The first way in which cognitive inflexibility may emerge in our task is as overall degradation in performance when switching demands are high compared to when they are low. This could manifest as a shift in performance as well as control settings on Varying relative to Fixed blocks, irrespective of the current performance goal. However, at odds with this prediction, at the group level, if anything, we observe performance to instead be better not worse during Varying blocks, reflected in faster (*b*=−8.51; 95% CrI [−21.08, 3.13]; *p*_*b*>0_=0.08) and more accurate (*b*=−0.09; 95% CrI [−0.18, −0.01]; *p*_*b*>0_=0.02) responding, and higher levels of both drift rate (*b*=−0.06; 95% CrI [−0.12, −0.01]; *p*_*b*>0_=0.02) and threshold (*b*=−0.02; 95% CrI [−0.06, 0.01]; *p*_*b*>0_=0.07l; Fig. 2D and Table S1). Nevertheless, these group-level effects could mask differences across our age range, including whereby the effect reverses to reveal switch-related performance decrements in older adults. We did not, however, find that this was the case. Increasing age actually led to even faster performance in varying relative to fixed blocks (*b*=−42.34; 95% CrI [−90.44, 9.22]; *p*_*b*>0_=0.05) and didn’t change differences in accuracy (*b*=−0.19; 95% CrI [−0.57, 0.21]; *p*_*b>*0_=0.18), nor did it predict differences in control configuration towards greater decreases in control on Varying relative to Fixed blocks (drift rate: *b*=−0.12; 95% CrI [−0.38, 0.12]; *p*_*b*>0_=0.19; threshold: *b*=−0.09; 95% CrI [−0.23, 0.05]; *p*_*b*>0_=0.12; Fig. 2C and Table S1). We next compared directly tested how these control configurations changed with age by calculating subject-level changes between Fixed and Varying blocks using a signed distance metric (positive for higher control intensity in Fixed vs. Varying blocks and negative otherwise; see Methods for details). Using this index of *switching* flexibility, we found a small block-related increase in control in Varying blocks that was not statistically significant (*b*=−0.03; 95% CrI [−0.08, 0.02]; *p*_*b*>0_=0.09; Fig. 2D and Fig. S2B).

The second way in which cognitive inflexibility could emerge in our task is through limitations in the extent to which older adults adjust their performance under the two goals. Older adults may be more rigid in their approach to performing the task, for instance by prioritizing achieving the Accuracy goal and therefore not adjusting to the Speed goal to the extent that younger adults do. This form of (in)flexibility can be indexed by the difference between performance (Fig. 1C), and corresponding control configurations (Fig., 1D) achieved under the two goals (orange vs. blue). We found that this form of flexibility was preserved with age as older adults maintained the same distance in their performance (RTs: *b*=37.08; 95% CrI [−48.05, 127.46]; *p*_*b*<0_=0.21; accuracy: *b*=−0.50; 95% CrI [−1.35, 0.34]; *p*_*b>*0_=0.11) or control configurations (drift: *b*=0.04; 95% CrI [−0.28, 0.39]; *p*_*b*<0_=0.42; threshold: *b*=−0.14; 95% CrI [−0.34, 0.04]; *p*_*b*>0_=0.08) under the two goals (Fig. 2A and Table S1). To more directly index how the distances between two control configurations changed with age, we extracted subject-level control configurations for the two goals and computed Euclidian distances for each subject (see Methods for details). This index of *configural flexibility* did not increase with age (*b*=−0.01; 95% CrI [−0.08, 0.07]; *p*_*b*>0_=0.44; Fig. 2B, Fig. S2A, and Table S1). In other words, participants converged on Speed and Accuracy configurations that were roughly equidistant from each other across age (despite the overall targets being offset from one another – biased towards higher thresholds – with age).

In stark contrast to the measures of flexibility above, we find that significant age-related differences emerge when examining indices of the time it takes to transition between control configurations. Specifically, we found that older adults undershoot target control configurations to a much greater degree than younger adults, despite traversing similar distances as young adults to adjust control configurations in accordance with a new goal. This is first reflected in their behavior, where we found that with increasing age the difference between speed and accuracy goals in varying compared to fixed blocks was increased (RTs: *b*=63.57; 95% CrI [−19.41, 140.28]; *p*_*b*<0_=0.06; accuracy: *b*=0.86; 95% CrI [0.13, 1.63]; *p*_*b*<0_=0.01). This is robustly reflected in the relative configurations of drift and threshold for varying compared to fixed blocks (Fig. 2E and Table S1), which fall closer to the center of the two target configurations (and further from the respective targets) for older relative to younger adults (drift rate: *b*=0.64; 95% CrI [0.22, 0.99]; *p*_*b*<0_=0.00; threshold: *b*=0.26; 95% CrI [0.07, 0.43]; *p*_*b*<0_=0.00). To quantify this difference, we computed subject-level changes in Euclidian distances between Speed and Accuracy goals in Fixed and Varying blocks. Using this index of *adjustment speed* we find that the amount of undershooting of control configurations in Varying relative to Fixed blocks increased significantly with age (*b*=0.08; 95% CrI [0.02, 0.14]; *p*_*b*<0_=0.00; Fig. 2F and Fig. S2C), suggesting that older adults take longer to transition from one control configuration to another.

Finally, while all the foregoing analyses tested for linear effects of age, we performed follow-up analyses to test for nonlinear effects (Table S2). Overall, we find that the linear effects reported above do not change qualitatively when including quadratic effects of age in the same regressions (for reaction times, accuracy, and computational model parameters). Notably, the only key change observed in these analyses is that monotonic reductions in adjustment speed with age are further enhanced when allowing for this nonlinearity.

## General Discussion

Achieving the various goals we face on a daily basis requires the ability to flexibly adjust how we perform a task depending on our current goals. The ability to rapidly adjust cognitive control configurations in accordance with our current goal is a critical feature of cognitive flexibility. While aging is known to be associated with diminished flexibility (Cepeda et al., 2001; Wasylyshyn et al., 2011; Chen & Hsieh, 2023), it remains unknown to what extent this can be attributed to changes in one’s ability to adjust control states; in their ability to maintain similar levels of control under changing goals; and/or in the time it takes them to adjust their control configuration to meet a new goal. By leveraging experimental and computational tools that we recently developed to disentangle each of these elements of cognitive flexibility (Grahek et al., 2025), we were able to test these each of the corresponding accounts of age-related changes in cognitive flexibility. Our results reveal a striking dissociation: whereas older adults are able to adjust their control configuration to a similar extent as younger adults, and to maintain similar overall levels of control in the face of changing goals, the speed with which they adjust their control is significantly slowed.

With increasing age, performance on cognitive tasks undergoes a number of significant changes. Relative to younger adults, older adults are typically slower to respond, but they tend to maintain their accuracy or even increase it. Computational models of response selection have uncovered age-related changes in control configurations that produce such changes: older adults exhibit larger levels of response caution (Theisen et al., 2021; von Krause et al., 2022). However, such results are commonly observed in experiments that employ a single performance goal (e.g., perform the task as quickly and accurately as you can). Here, we demonstrate that both younger and older adults modulate their performance and control configurations to a similar degree when asked to perform a control demanding task (Stroop) under two goals: focusing on accuracy or on speed. Thus, our results suggest that *configural flexibility*, the ability to implement different control configurations in accordance with the current goal, is maintained across age. Furthermore, by comparing performance in blocks that demand frequent control adjustments (varying goals blocks) to performance in blocks requiring a single control configuration (fixed goal blocks), we show that older relative to younger adults don’t display diminished control when having to frequently change goals (*switching flexibility*). Thus, despite evidence that older adults exhibit worse performance when required to switch between tasks (Cepeda et al., 2001; Chen & Hsieh, 2023; Mayr, 2001; Wasylyshyn et al., 2011), they seem to exhibit similar levels of performance when performing the same task under different goals.

Our key findings is that increasing age is related to slower control adjustments. Comparing control configurations in blocks that require frequent control adjustments to blocks that do not allow us to obtain a measure of the time it takes for people to adjust control states. Our previous work shows that when people’s performance goals change (e.g., from speed to accuracy focus) they gradually increase levels of attentional focus (drift rate parameter) and response caution (threshold parameter; Grahek et al., 2025). Due to the gradual movement from the current to the target state required by the new goal, people often undershoot that target state. Thus, the reduction in control configuration distance in varying relative to fixed blocks serves as the index of speed of control adjustment. This reduction is smaller in people with fast adjustments because they reach the target state faster. Critically, this speed is substantially reduced with age, despite preserved distance between control configurations across age. This result suggests that while older adults can reach different control configurations, they often fail to do so due to the time it takes them to reach the configuration appropriate for the current goal.

Our finding that the speed of control adjustments substantially reduces with age emphasizes the importance of the dynamical view of cognitive flexibility. Under this view, cognitive flexibility depends on the ability to transition between different cognitive and neural states when one’s tasks or goals change (Braun et al., 2015; Ueltzhöffer et al., 2015; Kim & Bassett, 2019; Musslick & Bizyaeva, 2024). There is already some evidence from the task switching literature suggesting that reduced speed of transitioning between task sets limits cognitive flexibility in older adults (Steyvers et al., 2019). At the neural level, older adults exhibit impaired transitions between neural states related to different tasks (Hakun et al., 2015; McDonald et al., 2018; Ezaki et al., 2018; Goodman et al., 2023), as well as slower dynamical transitions in resting state brain activity (Ezaki et al., 2018) and reduced modulation of activity in cognitive control regions in response to changing task demands (Kennedy et al., 2015). By leveraging model-based analyses of a task designed to selectively modulate control over different timescales, our current work provides a more direct measure of age-related changes in control configurations under varying demands on cognitive flexibility. While the neural mechanisms of observed changes in control dynamics are out of scope of the current paper, one prominent candidate is the global age-related change in neuronal gain. Reduction in gain will translate into slower control adjustments in our computational model of control dynamics and induce larger adjustment costs (Grahek et al., 2025). Norepinephrine (NE) is a critical neuromodulator which changes neuronal gain (inhibition-excitation balance) across cortex, regulating neural dynamics with higher gain leading to faster dynamics (Aston-Jones & Cohen, 2005; Poe et al., 2020; Wainstein et al., 2022). Locus coeruleus (LC) is a brainstem nucleus responsible for changing cortical levels of norepinephrine, and thus playing an important role in behavioral flexibility and cognitive control (Aston-Jones et al., 2007; Dahl et al., 2022; Grueschow et al., 2020; Kane et al., 2017; McBurney-Lin et al., 2022; Unsworth & Robison, 2017). Critically, the LC-NE system undergoes profound changes in both healthy and disordered aging (Clewett et al., 2016; Elman et al., 2017; Mather & Harley, 2016; Poe et al., 2020). Investigating the connection between neuronal gain, neuromodulators, and control dynamics across age could elucidate the mechanisms of reduced cognitive flexibility across age.

In this work, we used a novel task paradigm to systematically study different possible causes of cognitive inflexibility in older adulthood. While previous work has focused on deficits in switching between different tasks (Cepeda et al., 2001; Goodman et al., 2023; Heckner et al., 2021; Steyvers et al., 2019), we have demonstrated that cognitive inflexibility can be caused by slow adjustments of control states within a single task under different goals. Measuring continuous cognitive control states that determine how people process information and select actions enabled us to precisely measure age-related changes in control dynamics. This approach allowed us to test different possible causes of cognitive inflexibility and demonstrate that increasing age is uniquely related to substantial decreases in the speed of adjusting control signals. In this way we demonstrate that increasing age is related to substantial changes in control dynamics which limit people’s ability to adjust their performance in accordance with the changes in their goals and/or other cognitive demands brought on by their environment.

## Methods

### Participants

We recruited participants from Prolific. All participants had normal or corrected-to-normal vision, were fluent English speakers, and resided in United States. Research protocol was approved by Brown University’s Institutional Review Board. The study took approximately 1h to complete, and participants were compensated with a fixed rate of $8 per hour. We recruited a total of 248 who completed the study, out of which 3 participants did not pass the attention checks. The final sample included 245 participants (98 males, 146 females, 1 without reported gender; age range = 19-88; median age = 53).

### Task Design

Participants performed a Stroop task in which their goal was to respond to the ink color while ignoring a color word. They pressed one of 4 keys that corresponded to one of the 4 possible colors (blue, red, yellow, or green). Stroop trials were created with equal chance of being either congruent (matching ink color and color word) or incongruent (different ink color and color word). Ink and word colors were randomized under the constraint that both the color word and the ink color did not repeat in two consecutive trials. Participants performed an interval-based version of the Stroop task in which they could complete as many trials as they wished within a fixed time interval (randomly sampled to be 8, 9, 10, 11, or 12 seconds long). In previous work we demonstrated that this version of the task is suitable for measuring adjustments in drift rates and thresholds induced by incentives (Grahek et al., 2022; Leng et al., 2021) or performance goals (Grahek et al., 2025). Before each interval participants saw a cue (1.5s) informing them that they should perform the task as quickly (speed goal) or as accurately (accuracy goal) as they could (Forstmann et al., 2008; Ratcliff & Rouder, 1998). After each interval, they received feedback (1.5 s) informing them about the number of correct responses they made during the interval. Participants performed the task in 4 blocks consisting out of 20 intervals each. Critically, half of the blocks had a fixed goal (e.g., speed goal only), while half of the blocks had goals randomly switching between speed and accuracy. While fixed blocks required a constant control state, varying blocks demanded frequent control adjustments, inducing control adjustment costs. Prior to the start of each blocks participants were informed of the block type, and block order was randomized for each participant. Before starting the main task, participants performed several practice blocks to familiarize themselves with the task. They first learned the color-button mapping, then performed the Stroop task, and then practiced doing the task in the speed, accuracy, and varying goals conditions. The task was coded in Psiturk (Gureckis et al., 2016) Participants did the task on their own computers and having a keyboard was required.

### Statistical Analyses and Drift Diffusion Modeling

#### Reaction Times and Accuracy

We fitted Bayesian multilevel regressions to predict reaction times (Ex-gaussian likelihood) and accuracy (Bernoulli likelihood). Goal type (speed vs. accuracy), Block type (fixed vs. varying), linear effect of Age, quadratic effect of age, and interactions between these variables were used as predictors in both models. We controlled for the effects of congruency, interval length, and the time spent in the interval. Congruency, Goal type, Block type and Goal-Block interactions were also entered as random effects, allowing them to vary across subjects. When analyzing reaction times, we included only the correct responses, and excluded reaction times shorter than 250ms and longer than 3000ms. Regression models were fitted in R using the *brms* (Bürkner, 2017; Nalborczyk et al., 2019) package which relies on *Stan* (Carpenter et al., 2017) to implement Markov Chain Monte Carlo (MCMC) algorithm and estimate posterior distributions of parameters. All models included default uniform priors. To estimate model parameters, we ran four MCMC chains with 20000 iterations per simulation, discarding the first 19000 chains (warmup). We confirmed convergence by examining trace plots, autocorrelation, variance between and within chains (Gelman & Rubin, 1992), and posterior predictive checks. For each parameter, we report the mean of the posterior distribution (*b*) and the credible intervals (95% CrI). We also report the proportion of the posterior samples on one side of 0 (e.g., p_*b* < 0_ = 0.01) which represents the probability that given parameter estimate is below 0. Raw data and analysis scripts are available on GitHub (https://github.com/igrahek/ControlFlexibilityAging.git).

#### Drift Diffusion Modeling

We jointly modeled reaction times and accuracy using a hierarchical Bayesian Drift Diffusion Model (Wiecki et al., 2013). We used the same exclusion criteria for reaction times as reported for the behavioral analyses above. Furthermore, we only included trials on which the incorrect response corresponded to the color word and thus driven by the automatic prepotent response to the word. We did this to exclude errors which are likely driven by random responding (Grahek et al., 2022, 2025; Leng et al., 2021). Following our previous work (Grahek et al., 2022, 2025), as predictors for drift-rate we used Goal type, Block type, their interaction, the linear and the quadratic effects of Age, the interactions between the Age variables with Goal type and Goal-Block interaction, Congruency (congruent vs. incongruent trials). Goal type, Block type, their interaction, and Congruency were also entered as random effects. We used the same predictors for threshold, but without the effect of Congruency and adding the effect of the amount of time elapsed within the interval. We also included the group and random effects of congruency on the starting point bias parameter, and the effect of Goal type on the non-decision time. Using the HSSM package in Python (Fengler et al., in preparation), we ran 4 MCMC chains (3000 iterations per simulation; 2000 warmup). As for the regression models described in the previous section, we report the mean of the posterior distribution, the credible intervals, and the probability that the parameter of interest is below 0.

#### Flexibility metrics analyses

The switching flexibility metric measures whether control intensity in varying relative to fixed blocks was reduced due to frequent switches between performance goals. To index this, we first obtained control configurations (drift rate and threshold values) from the fitted Drift Diffusion model by taking the means of the condition-wise posterior distributions. Next, we projected these points onto the unit diagonal vector, allowing us to ask whether the overall control intensity has decreased or increased when comparing fixed to varying blocks. In this way we obtained a signed distance metric, with positive values indicating increased control signals in fixed relative to varying blocks, and negative values indicating the opposite.

The configural flexibility metric measures the distance between control configurations under the Speed and Accuracy performance goals respectively. This metric was obtained by calculating the Euclidian distances between the two control configurations obtained from the respective posterior distributions. This metric is not signed as the index of configural flexibility is the distance between the two conditions regardless of the relative positions of the two goals in the drift rate – threshold space.

The metric of adjustment speed was obtained by first calculating the Euclidian distance between the Speed and Accuracy configurations in Fixed and Varying blocks respectively. Next, we subtracted the distance in Fixed from the distance in Varying blocks to calculate the speed metric. Larger values of this metric indicate increased reduction in distances between Fixed and Varying blocks, suggestive of slower speed of travel between the target configurations.

To analyze how the three flexibility metrics changed with age, we extracted the subject-level posterior estimates of each of the four conditions from the fitted Drift Diffusion model. Next, we calculated subject-level estimates of all three metrics. Since there are individual differences in the distances between condition-wise configurations, when calculating the adjustment speed metric we first min-max range normalized posterior samples across all conditions for each subject. To test how each of the metrics change with age, we fit three Bayesian linear regression models to predict each of the three metrics by participant’s age. Similarly to the procedure described in more detail above, we estimated model parameters running four MCMC chains with 4000 iterations per simulation, discarding the first 2000 chains (warmup).

## Acknowledgements

NIMH R01MH124849 (A.S.), NSF Career Award 204611 (A.S.), and NIH Award 1S10OD02518 (A.S.). We thank Meriel Doyle for her help during data collection.

## Supplementary materials

**Figure S1.**
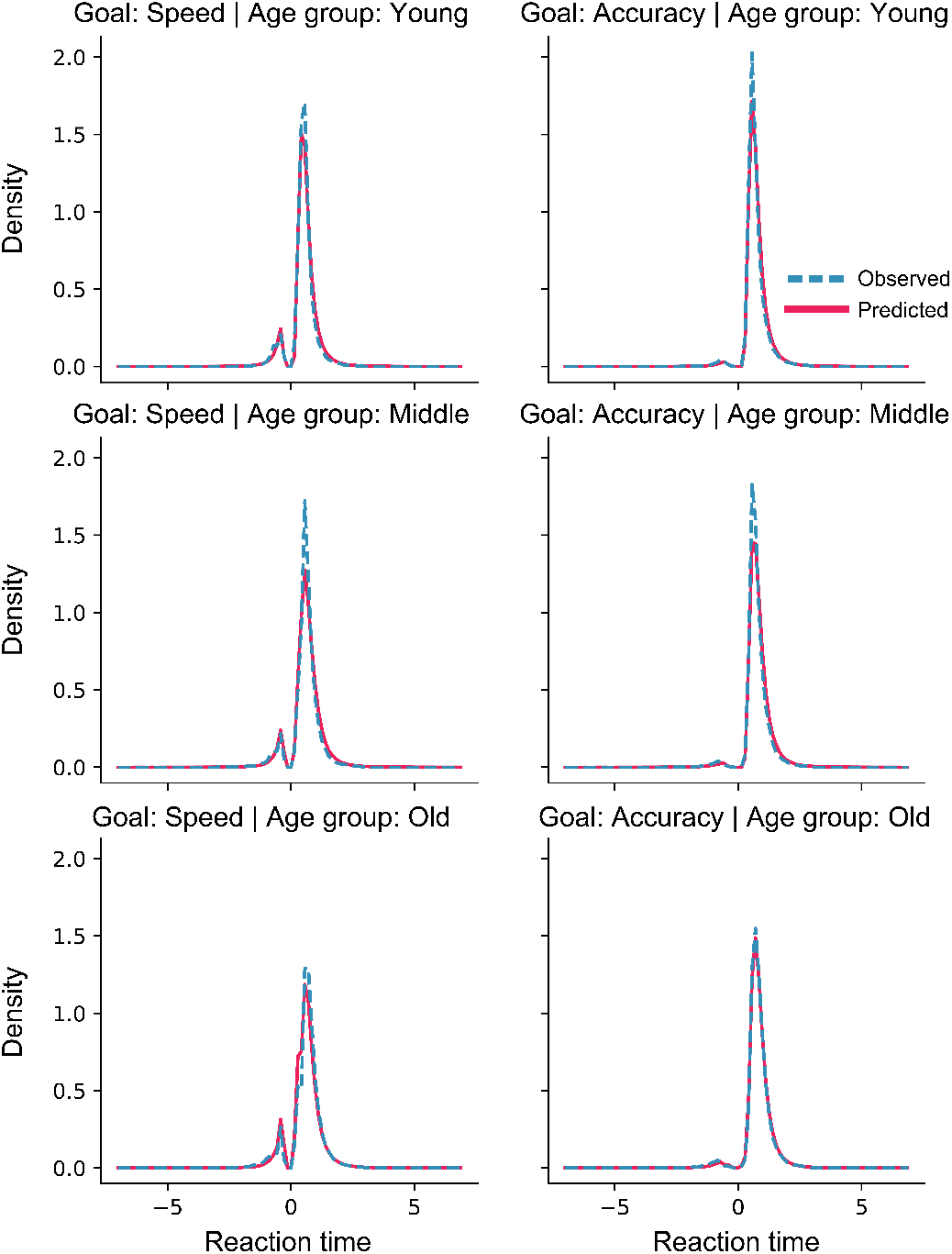
Posterior predictive checks for the Diffusion model. Distributions of simulated (red) and observed (blue) reaction times for correct (positive reaction time) and incorrect responses (negative reaction time). The model captures the observed data well in both speed and accuracy goal conditions (columns) and across age terciles (rows).

**Table S1.**
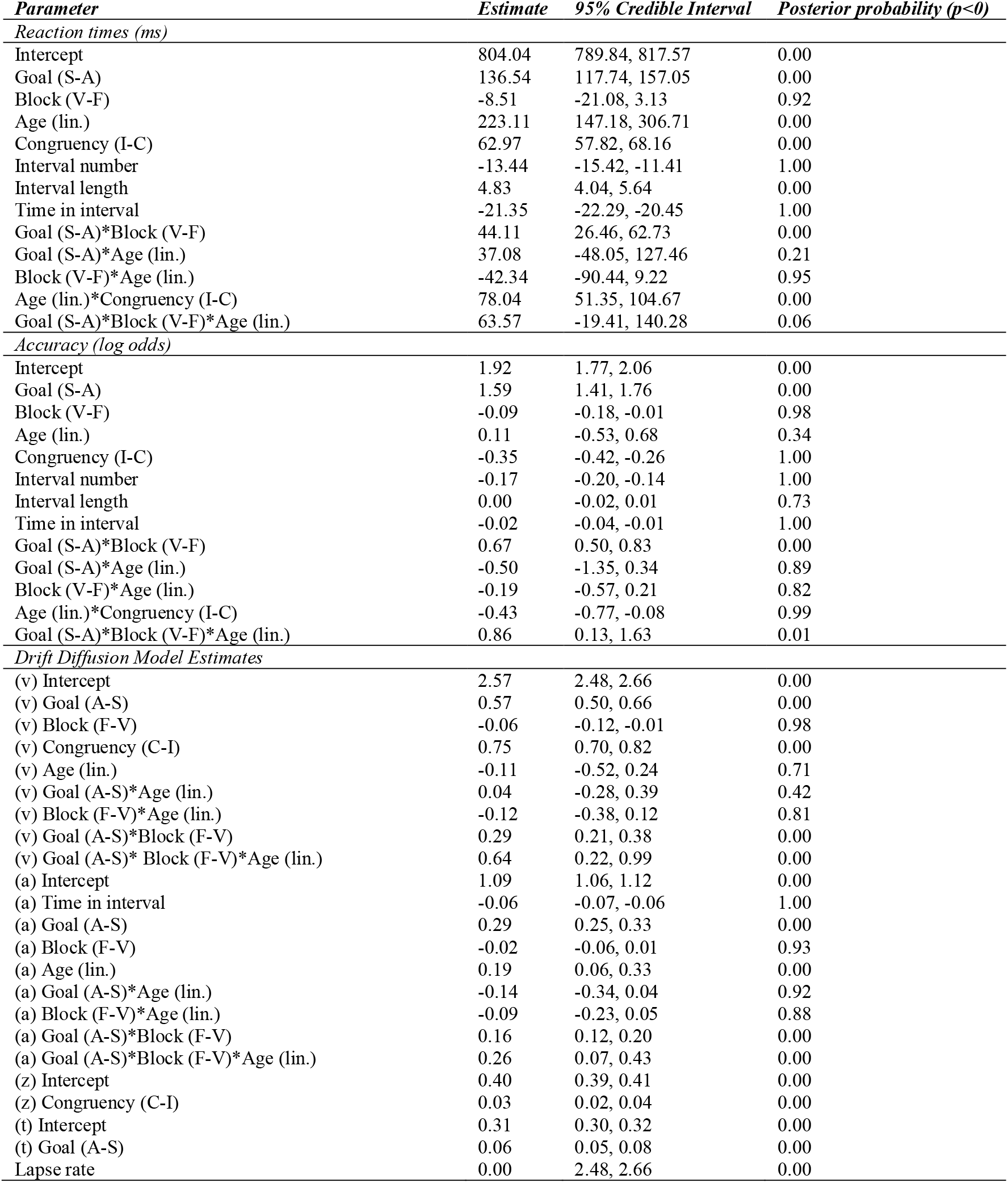
Parameter estimates for hierarchical regressions with the linear (lin.) effect of age predicting behavioral performance (reaction times and accuracy) and parameter estimates of the Drift Diffusion Model (a – threshold; v – drift rate; t – non-decision time; z – bias).

**Table S2.**
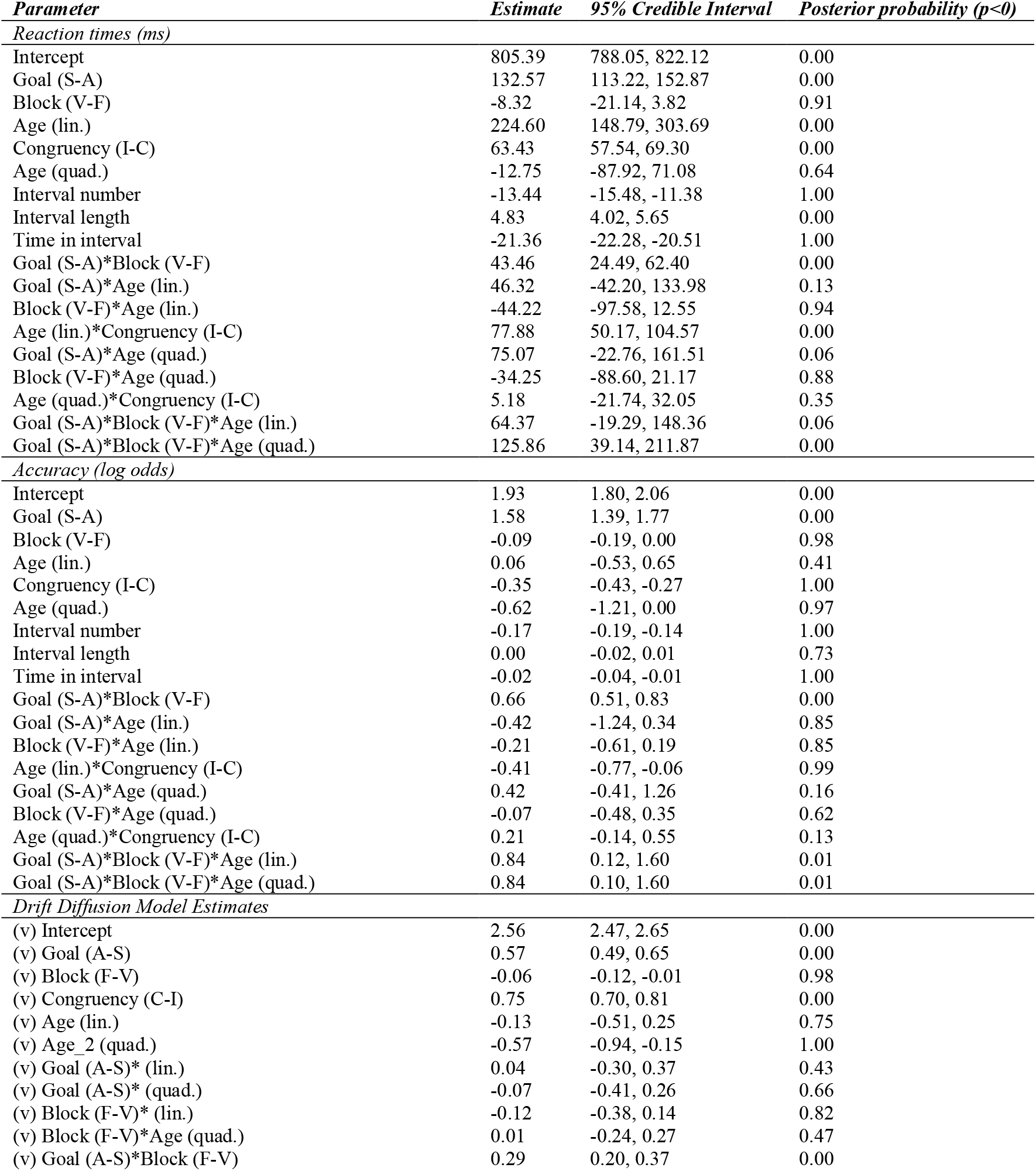
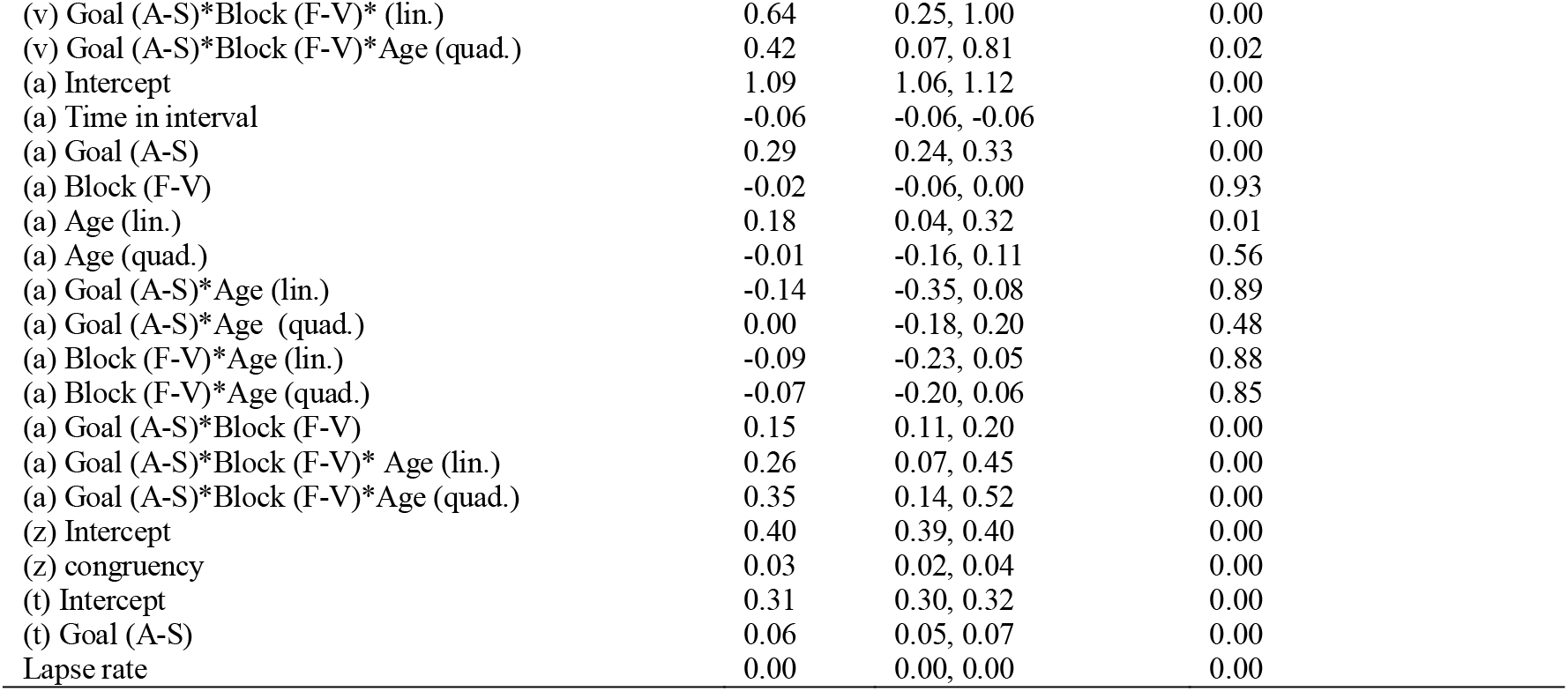
Parameter estimates for hierarchical regressions with the linear (lin.) and quadratic (quad.) effects of age predicting behavioral performance (reaction times and accuracy) and parameter estimates of the Drift Diffusion Model (a – threshold; v – drift rate; t – non-decision time; z – bias).

| *Parameter* | *Estimate* | *95% Credible Interval* | *Posterior probability (p<0)* |
| --- | --- | --- | --- |
| *Reaction times (ms)* |  |  |  |
| Intercept | 805.39 | 788.05, 822.12 | 0.00 |
| Goal (S-A) | 132.57 | 113.22, 152.87 | 0.00 |
| Block (V-F) | -8.32 | -21.14, 3.82 | 0.91 |
| Age (lin.) | 224.60 | 148.79, 303.69 | 0.00 |
| Congruency (I-C) | 63.43 | 57.54, 69.30 | 0.00 |
| Age (quad.) | -12.75 | -87.92, 71.08 | 0.64 |
| Interval number | -13.44 | -15.48, -11.38 | 1.00 |
| Interval length | 4.83 | 4.02, 5.65 | 0.00 |
| Time in interval | -21.36 | -22.28, -20.51 | 1.00 |
| Goal (S-A)*Block (V-F) | 43.46 | 24.49, 62.40 | 0.00 |
| Goal (S-A)*Age (lin.) | 46.32 | -42.20, 133.98 | 0.13 |
| Block (V-F)*Age (lin.) | -44.22 | -97.58, 12.55 | 0.94 |
| Age (lin.)*Congruency (I-C) | 77.88 | 50.17, 104.57 | 0.00 |
| Goal (S-A)*Age (quad.) | 75.07 | -22.76, 161.51 | 0.06 |
| Block (V-F)*Age (quad.) | -34.25 | -88.60, 21.17 | 0.88 |
| Age (quad.)*Congruency (I-C) | 5.18 | -21.74, 32.05 | 0.35 |
| Goal (S-A)*Block (V-F)*Age (lin.) | 64.37 | -19.29, 148.36 | 0.06 |
| Goal (S-A)*Block (V-F)*Age (quad.) | 125.86 | 39.14, 211.87 | 0.00 |
| *Accuracy (log odds)* |  |  |  |
| Intercept | 1.93 | 1.80, 2.06 | 0.00 |
| Goal (S-A) | 1.58 | 1.39, 1.77 | 0.00 |
| Block (V-F) | -0.09 | -0.19, 0.00 | 0.98 |
| Age (lin.) | 0.06 | -0.53, 0.65 | 0.41 |
| Congruency (I-C) | -0.35 | -0.43, -0.27 | 1.00 |
| Age (quad.) | -0.62 | -1.21, 0.00 | 0.97 |
| Interval number | -0.17 | -0.19, -0.14 | 1.00 |
| Interval length | 0.00 | -0.02, 0.01 | 0.73 |
| Time in interval | -0.02 | -0.04, -0.01 | 1.00 |
| Goal (S-A)*Block (V-F) | 0.66 | 0.51, 0.83 | 0.00 |
| Goal (S-A)*Age (lin.) | -0.42 | -1.24, 0.34 | 0.85 |
| Block (V-F)*Age (lin.) | -0.21 | -0.61, 0.19 | 0.85 |
| Age (lin.)*Congruency (I-C) | -0.41 | -0.77, -0.06 | 0.99 |
| Goal (S-A)*Age (quad.) | 0.42 | -0.41, 1.26 | 0.16 |
| Block (V-F)*Age (quad.) | -0.07 | -0.48, 0.35 | 0.62 |
| Age (quad.)*Congruency (I-C) | 0.21 | -0.14, 0.55 | 0.13 |
| Goal (S-A)*Block (V-F)*Age (lin.) | 0.84 | 0.12, 1.60 | 0.01 |
| Goal (S-A)*Block (V-F)*Age (quad.) | 0.84 | 0.10, 1.60 | 0.01 |
| *Drift Diffusion Model Estimates* |  |  |  |
| (v) Intercept | 2.56 | 2.47, 2.65 | 0.00 |
| (v) Goal (A-S) | 0.57 | 0.49, 0.65 | 0.00 |
| (v) Block (F-V) | -0.06 | -0.12, -0.01 | 0.98 |
| (v) Congruency (C-I) | 0.75 | 0.70, 0.81 | 0.00 |
| (v) Age (lin.) | -0.13 | -0.51, 0.25 | 0.75 |
| (v) Age_2 (quad.) | -0.57 | -0.94, -0.15 | 1.00 |
| (v) Goal (A-S)* (lin.) | 0.04 | -0.30, 0.37 | 0.43 |
| (v) Goal (A-S)* (quad.) | -0.07 | -0.41, 0.26 | 0.66 |
| (v) Block (F-V)* (lin.) | -0.12 | -0.38, 0.14 | 0.82 |
| (v) Block (F-V)*Age (quad.) | 0.01 | -0.24, 0.27 | 0.47 |
| (v) Goal (A-S)*Block (F-V) | 0.29 | 0.20, 0.37 | 0.00 |
| (v) Goal (A-S)*Block (F-V)* (lin.) | 0.64 | 0.25, 1.00 | 0.00 |
| (v) Goal (A-S)*Block (F-V)*Age (quad.) | 0.42 | 0.07, 0.81 | 0.02 |
| (a) Intercept | 1.09 | 1.06, 1.12 | 0.00 |
| (a) Time in interval | -0.06 | -0.06, -0.06 | 1.00 |
| (a) Goal (A-S) | 0.29 | 0.24, 0.33 | 0.00 |
| (a) Block (F-V) | -0.02 | -0.06, 0.00 | 0.93 |
| (a) Age (lin.) | 0.18 | 0.04, 0.32 | 0.01 |
| (a) Age (quad.) | -0.01 | -0.16, 0.11 | 0.56 |
| (a) Goal (A-S)*Age (lin.) | -0.14 | -0.35, 0.08 | 0.89 |
| (a) Goal (A-S)*Age (quad.) | 0.00 | -0.18, 0.20 | 0.48 |
| (a) Block (F-V)*Age (lin.) | -0.09 | -0.23, 0.05 | 0.88 |
| (a) Block (F-V)*Age (quad.) | -0.07 | -0.20, 0.06 | 0.85 |
| (a) Goal (A-S)*Block (F-V) | 0.15 | 0.11, 0.20 | 0.00 |
| (a) Goal (A-S)*Block (F-V)* Age (lin.) | 0.26 | 0.07, 0.45 | 0.00 |
| (a) Goal (A-S)*Block (F-V)*Age (quad.) | 0.35 | 0.14, 0.52 | 0.00 |
| (z) Intercept | 0.40 | 0.39, 0.40 | 0.00 |
| (z) congruency | 0.03 | 0.02, 0.04 | 0.00 |
| (t) Intercept | 0.31 | 0.30, 0.32 | 0.00 |
| (t) Goal (A-S) | 0.06 | 0.05, 0.07 | 0.00 |
| Lapse rate | 0.00 | 0.00, 0.00 | 0.00 |

**Figure S2.**
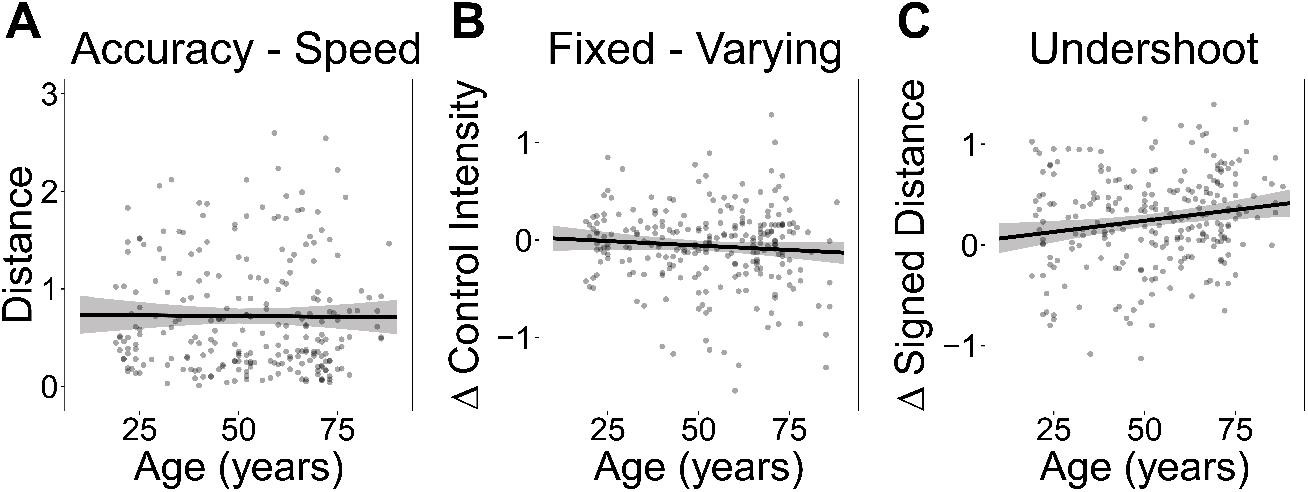
Subject-level model-based estimates of the influence of age on different flexibility metrics. **A**. Euclidian distance between speed and accuracy conditions across age. Distances are calculated by obtaining the means of the condition-level posteriors for each subject from the hierarchical Drift Diffusion model. Error bars represent 95% credible intervals **B**. Signed distance between the fixed and varying conditions across age. Euclidian distances are projected onto the unit vector along the main diagonal, allowing the distance to be negative if the Varying condition is above the Fixed condition along the diagonal. **C**. The level of undershooting the target control configuration in varying relative to fixed goal blocks across age. The undershoot metric for each subject is obtained as the reduction in Euclidian distance between the accuracy and the speed configurations in varying relative to fixed blocks.

